# Different Diets Modulate the Gut Microbiome Compositions and Promote the Health of *Apis mellifera*

**DOI:** 10.1101/2023.11.26.568759

**Authors:** Hyun Jee Kim, Abdulkadir Yusif Maigoro, Jeong Hyeon Lee, Olga Frunze, Mustafa Bilal, Hyung Wook Kwon

## Abstract

Honey bee (*Apis mellifera*) health is crucial for honey bee products, and effective pollination and is closely associated with gut bacteria. Various factors such as reduced habitat, temperature, disease, and diet affect the health of honey bees, by disturbing the homeostasis of the gut microbiota. In this study, high-throughput 16S rRNA gene sequencing was used to analyze the gut microbiota of *Apis mellifera* subjected to seven different diets. The identified microbiota in the *Apis mellifera* gut from all the diets consisted of *Lactobacillus* (62%), followed by *Rhizobiaceae* (21%), *Snodgrassella* (4%), and *Erwiniaceae* (4%) among other 33 genera. Based on diet types, *Lactobacillus* a lactic acid bacteria (LAB), dominates the microbiota with the highest relative abundance in AIGT+SAC (91%), AIGT+Soytide (88%), and AIGT+Apple juice (69%) diet groups. *Bifidobacterium* and *Commensalibacter* appeared as the second most abundant genera in AIGT+SAC and AIGT+Soytide diet groups, respectively. These bacteria are important markers for honey bee health. Considering the importance of these diets in shaping their host microbiome into a healthy status. Individual honey bee health (IHH) was observed to validate the quality and correlation between the microbiota and honey bee health. The results were consistent, indicating that *Apis mellifera* fed on AIGT+Soytide and AIGT+SAC diet showed the highest health expression level of vitellogenin. The group with 60%Syrup possessing *Rhizobiaceae* as the dominant taxa showed poor health status. This finding paved the way for establishing a link between gut microbiota and IHH under different diets.

## 1.0 Introduction

Honey bees (*Apis mellifera*) are crop pollinators of economic importance, widely used in agriculture and food production [1]. The gut of honey bees is occupied with a large portion of microbes which can impact honey bee pollinators in several ways, such as nutrition, development, and defense against diseases [2]. The role of gut microbiome in animal health has been established ranging from mammals to insects [3–5]. Their importance in honey bee health and diseases has recently been emphasized [6]. Unlike higher animals, the honey bee core microbiome occupies about 95% of all gut bacteria [7]. The core microbiome are grouped into 9 core bacterial species belong to Alphaproteobacteria, Betaprotobacteria, Gammaproteobacteria, Firmicutes, and Actinobacteria [8,9].

Further, the honey bee microbiota compositions keep on changing according to development stage, diet, and environment [10,11]. These affect honey bee health status by either strengthening or weakening their immunity by subjecting them to disease susceptibility [12,13]. Diseases and pathogens affect not only honey bee health but also microbiome dysbiosis [7]. A high abundance of γ-proteobacterium was observed in the gut microbial community of honey bees undergoing colony collapse disorder (CCD) [7,8]. An elevated presence of γ-proteobacterium could either have detrimental effects on honey bee health or play a beneficial role in enhancing the resilience of insect hosts against gastrointestinal threats [14]. On the other hand, a high abundance of *Lactobacillus*, and Actinobacteria (*Bifidobacterium*) were observed in diseased honey bees [15]. These two symbiotic microbes can adapt to various microenvironments and play a protective role against microbial invasion in honey bees [16,17]. This suggests that an elevated amount of *Lactobacillus* and *Bifidobacterium* were beneficial to improve host immunity to honey bee damage. Similarly, the abundance of these microbiome correlates with the significant enrichment of carbohydrate metabolism in the diseased honey bee, which reflects a unique adaptation of gut microorganisms to diet [18]. Diets have effects on shaping the gut microbiota in insect species [19,20]. In contrast with genetic factors, diet outweighs genetics in shaping gut microbiota in Asian honey bees [21].

Even though it is affected by season, temperature, development stage, and diseases, among other factors, the response of honey bee gut microbiota to different diets remains a topic of discussion. Added to the importance of diet in shaping the honey bee gut microbiota for health stability and immune boosting against diseases and pathogens. We developed different diets enriched in various nutrient compositions and fed them to the honey bees. Apart from checking the honey bee microbiome changes from different diets, honey bee individual performance was also evaluated. These include their diet consumption, protein content in the head, and vitellogenin (Vg) expression level. In this study, a correlation between microbiome, and honey bee individual performance was determined by using high-throughput 16S rRNA gene sequencing.

## 2.0 Material and Methods

### 2.1 Honey bee sampling

The cage experiment was conducted for five days at 33±2°C and 55±5% relative humidity conditions in an incubator. We chose a health colony from the Incheon National University Apiary then, took one capped brood frame and carried it to an incubator. Each cage (17 × 14.5 × 9.5 cm) was utilized for every twenty honey bees that emerged within 24 hours from the carried frame.

We developed pollen substitute diets which are named AIGT+Soytide, AIGT+Apple juice, AIGT+ SAC, and AIGT+ MAC. The specific composition of diets was listed in the previous paper [22]. We formulated AIGT which included TestA (A) as the first development diet, inosine-5’-monophosphate (I), guanosine-5’-monophosphate (G), and tangerine juice (T). The inosine-5’-monophosphate (I) and guanosine-5’-monophosphate (G) were added to improve umami taste reactions and purchased from Sigma Aldrich (Milwaukee, USA) [23]. Soytide, Apple juice, Chlorella, and M are added functional additive components to each diet formula. Soytide (CJ Global Food and Bio Company) was incorporated as a functional additive because of its attributes as a fermented soybean rich in protein and with minimal anti-nutritional and allergenic properties, fostering both development and efficient nutrient utilization. Apple juice (Jaan Company) was added as a longevity support ingredient due to result in high longevity among adult codling moths, *Cydia pomonella*. Chlorella (Cheonil Herbal Medicine) was added to show the effect of colony formation and boosted Vg synthesis in honey bees. AIGT+SAC was formulated with AIGT, Soytide (S), apple juice (A), and chlorella (C). AIGT+ MAC was formulated with M which is a fermenet form of defatted soy flour instead of Soytide from AIGT+SAC. Megabee (Castle Dome Solutions, Helena, AR, USA) is a commercial diet that is a blend of plant-based proteins devoid of pollen as a positive control [24], Beebread and honey were used as a natural condition, and 60% Syrup used as a negative control. Three independent cages were used as replications for each diet, and 20g of each diet was provided with boiled water freely dispensed through a hole pierced in the cap of a 15ml tube using a pin. The tube was then placed upside down on the top of the cage.

### 2.2 DNA extraction

After five days of feeding, we sampled 10 live honey bees from each cage and placed them individually in 15ml tubes. The samples were stored in a liquid nitrogen container and kept at - 80°C until the experiment. Before dissection, honey bees underwent a 1-minute surface sterilization in 70% ethanol, followed by dissection in PBS. The whole gut of each of the three honey bees was used to extract genomic DNA from their respective tube. Under aseptic conditions, whole genome DNA was extracted using the Power soil kit (47014; QIAGEN, Hilden, Germany), according to the manufacturer’s instructions.

### 2.3 RNA Extraction and Quantitative PCR (qPCR)

Three honey bees were obtained from each diet sampling tube, three abdomens were pooled to extract RNA using a Qiagen RNeasy Mini Kit (#74104; Qiagen, Valencia, CA, USA). We analyzed Vg gene expression levels using quantitative PCR (qPCR) with cDNA templates from total RNA. For cDNA synthesis, 1 μg of total RNA was combined with oligo-dT and Invitrogen Superscript III enzyme. qPCR was performed on the Applied Biosystems StepOne Plus system with SYBR green qRT-PCR Master Mix from Fermentas. The cycling conditions included initial denaturation at 95 °C for 5 minutes, followed by 40 cycles of denaturation at 95 °C for 30 seconds, annealing at 60 °C for 30 seconds, and extension at 72 °C for 30 seconds. Details of qPCR primers are provided in (Table S1). Results were normalized to the control gene β-Actin using the 2-ΔΔCt method [25], and each biological replicate underwent technical triplicate testing.

### 2.4 Consumption

The quantity of diet consumption was determined by calculating the difference between 24 hours and dividing it by the number of live honey bees for that day. The daily consumption was then summed over the total experiment.

### 2.5 Soluble protein content

Quantitative measurement of water-soluble protein performed using Pierce BCA Protein Assay Kit (Thermo Scientific, Rockford, IL) [26]. The water-soluble protein content of the process was analyzed following the method with minor modifications, on SynergyTM HTX Multi-Mode Reader (BioTeK, USA) [27]. The water-soluble protein content was calculated as µg/ml in the head.

### 2.6 Sample preparation, 16s sequencing, and Taxonomic Analysis

For all samples, the National Instrumentation Center for Environmental Management (NICEM, www.nicem.snu.ac.kr), Republic of Korea, performed commercial PCR amplification, sample processing, and 16S rRNA gene sequencing. The samples were amplified using the KAPA HiFi HotStart ReadyMix (kk2601; Roche, Basel, Switzerland) and primers for the V3−V4 region of the 16S rRNA gene (Table S2). The following were the PCR conditions: 3 min at 96 °C, then 30 cycles of 30 s at 96 °C, 30 s at 55 °C, 30 s at 72 °C, and finally 5 min at 72 °C. All the PCR results were then performed on 1.2% agarose gels to determine band size and intensity. Ampure XP beads (A63882; Beckman, CA, USA) were used to purify amplified DNA from each sample. According to the content of DNA and molecular weight, samples were pooled in identical quantities and utilized to create Illumina DNA libraries. The libraries were then sequenced in Illumina MiSeq runs to obtain 2×300bp paired-end reads. The SILVA database v138.1 was used for taxonomic analysis. Protocol was adopted according to Lee et al [4].

### 2.7 Analysis in QIIME2

The sequencing data was analyzed using the Quantitative Insights into Microbial Ecology (QIIME2) pipeline [28]. The raw reads were denoised and trimmed using the DADA2 pipeline [29]. All the data were individually denoised before being merged for further analysis. The QIIME2 diversity plugin was used to construct the alpha and beta diversity index. The sampling depth was set at 3800. The bacterial diversity of honey bees from different diet groups was compared using a Shannon index. Using the QIIME2, the Shannon diversity index for honey bee samples was calculated using bacterial OTU count data. The Kruskal-Wallis H test was used to compare these results among developmental stages. To discover significant differences in their bacterial profiles, pairwise PERMANOVA was used with the ’beta-group-significance’ tool in QIIME2.

### 2.8 Bacterial profiles

At the class, family, and genus levels, read count and abundance data for bacterial OTUs were evaluated. Low abundance taxa with a value of less than one percent were categorized as “ETC” from the dataset. Then, the relative abundance for each bacterial OTU was calculated across all samples. The average and maximum abundance of bacterial genera in the dataset were used to identify abundant bacterial genera.

*2.9 Statistical Analysis*

The statistical analyses which Wilcoxon rank-sum test, PERMANOVA (Permutational multivariate analysis of variance), Kruskal-Wallis H test, Linear discriminant analysis Effect Size (LEfSe), and ANCOM analysis were performed using R version 4.1.0 in RStudio Version 1.4.1106 and QIIME2 [30]. We conducted data analysis using SPSS version 25 (IBM). To identify significant differences among means at a significance level of P < 0.05, we employed Duncan’s multiple range tests. The results are presented as mean values accompanied by their standard deviations (SD). Graphs were created using GraphPad Prism software (version 7.03).

## 3.0 Results

### 3.1 Reads Profiling For Microbial Community

Next-generation sequencing (NGS) returned an average of 71,394 quality trimmed reads with (∼4000bp maximum sequence depth). Out of 21 samples, the 60% Syrup diet has three samples with total non-chimeric reads of 34,059. AIGT+Apple juice diet has three samples with a total of 26,356 non-chimeric reads. Beebread and honey diet possessing three samples has a total number of 27,493 non-chimeric reads. The AIGT+MAC diet with three samples possesses a total number of 34,912 non-chimeric reads. Also, the Megabee diet has three samples with a total of 24,008 non-chimeric reads. The AIGT+SAC diet with three samples possesses a total number of 31,450 non-chimeric reads. However, the AIGT+SAC2 result shows a highly diverse result, and as such not considered for taxa relative abundance analysis. Finally, the AIGT+Soytide diet also has three samples with a total of 32,491 non-chimeric reads **(Table S3)**.

### 3.2 Gut Microbiota Diversity and Richness Between Different Diet Groups

#### 3.2.1 Alpha diversity

To investigate whether different diet compositions can affect the microbial community in *A. mellifera*, the Shannon index was evaluated. The alpha diversity was calculated which shows that there is slightly a considerable variation between the 60% Syrup diet and AIGT+Apple juice (**Figure 1**). Also between AIGT+Apple juice and Megabee. Similar variation was observed between AIGT+Apple juice and AIGT+SAC, as well as AIGT+Apple juice, and AIGT+Soytide diet groups respectively (**Figure 2**). The Kruskal-Wallis statistical analysis shows their respective P-value **(Table 1).** The Shannon index of all the significant values between the diet combinations falls within the P-values of 0.490 respectively. Other non-significant diet groups include the Beebread and honey diet from the Megabee diet with a P-value of 0.512, Megabee from the AIGT+SAC diet with a P-value of 0.275, and the AIGT+SAC diet from the AIGT+Soytide diet with a P*-*value of 0.512 **(Table 1)**. The principal coordinate analysis (PCoA) was further used to identify clustering patterns among the seven diet groups *viz* AIGT+Apple juice, Beebread and honey, AIGT+MAC, Megabee, AIGT+SAC, AIGT+Soytide, and 60% Syrup showed that the microbiome community structure of AIGT+Soytide and AIGT+Apple juice diets were separated from AIGT+MAC diets groups (**Figure 2**). Further, the 60% Syrup diet microbiome are in close relationship with a group of AIGT+MAC diet bee microbiome, which indicates that they have a close microbiome association, unlike the AIGT+SAC diet and Beebread and honey which have different microbiome communities. This data indicated that different diet combinations significantly affected the honey bee microbiome community structure under natural conditions.

**Figure 1.**
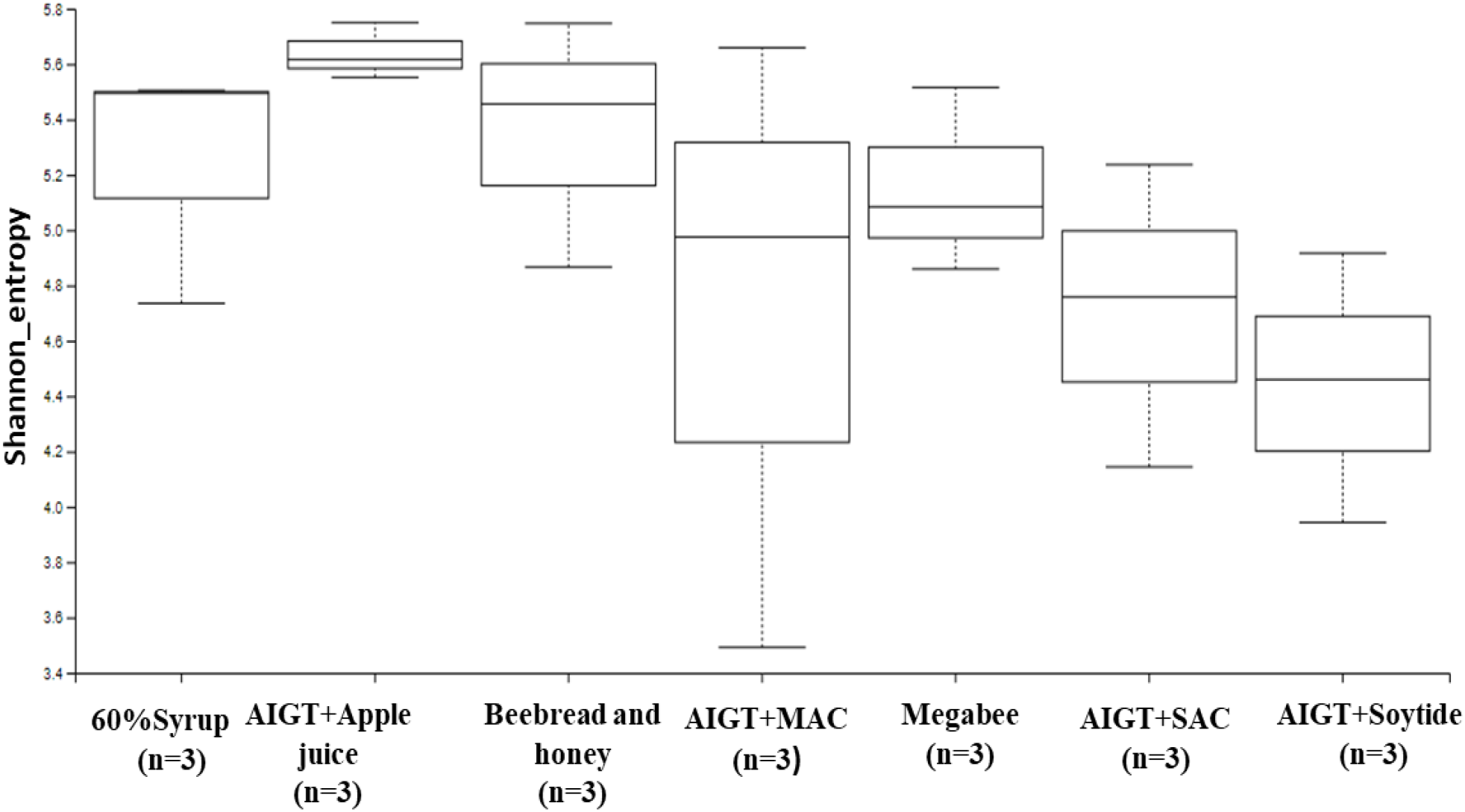
The Shannon entropy diversity index was performed using statistical analysis to measure the degree of randomness of the microbiome diversity within a sample based on species diversity and species richness of each diet group sample.

**Figure 2.**
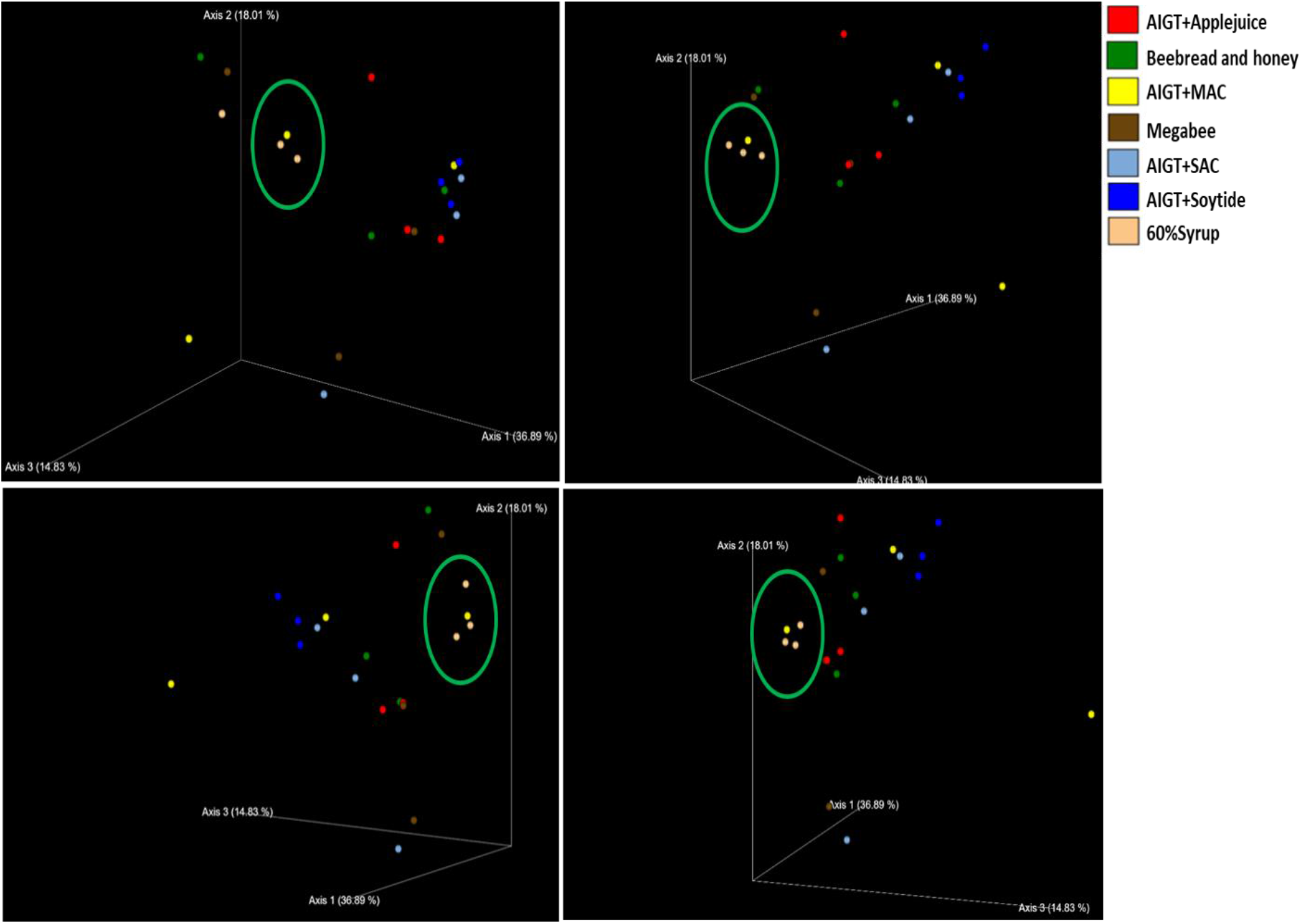
Score plot for principal coordinate analysis (PCoA) of the Bray-Curtis distances of the gut microbiome of *Apis mellifera*, using a multivariate analysis method. Data were expressed as the mean ± SEM (n = 7). This figure offers images that were shot from various angles for clear visualization. The PCoA plot was made based on the distance between different diet combinations based on their dissimilarities in terms of relative abundance (Bray-Curtis).

**Table 1.** Shannon index calculated P-values evaluated from the microbiome diversity within the respective diets. P-value < 0.05 shows significance.

| Group 1 | Group 2 | P-value | q-value |
| --- | --- | --- | --- |
| 60% Syrup (n=3) | AITG+Apple juice (n=3) | 0.049 | 0.260 |
| AITG+Apple juice (n=3) | Megabee (n=3) | 0.049 | 0.260 |
| AITG+Apple juice (n=3) | AITG+SAC (n=3) | 0.049 | 0.260 |
| AITG+Apple juice (n=3) | AITG+Soytide (n=3) | 0.049 | 0.260 |
| Beebread and honey (n=3) | Megabee (n=3) | 0.512 | 0.672 |
| Megabee (n=3) | AITG+SAC (n=3) | 0.275 | 0.481 |
| AITG+SAC (n=3) | AITG+Soytide (n=3) | 0.512 | 0.672 |

#### 3.2.2 Beta diversity

Beta diversity metrics were calculated between the samples of varying diet groups to test the distance relationship between the diet groups. Boxplots of unweighted UniFrac were constructed, incorporating distances in phylogenic trees among the diet group members in comparison with other groups. The result was similar to alpha diversity. We observed higher internal distances (mean: 60% Syrup, 0.32; AIGT+Apple juice, 0.51; Beebread and honey, 0.82; AIGT+MAC, 1.0; Megabee, 0.73; AIGT+SAC, 0.79; and AIGT+Soytide, 0.53) of diet members and the highest distance between the groups (mean: 60% Syrup-Beebread and honey, 0.7; 60% Syrup-AIGT+Soytide, 0.9; AIGT+Soytide-Megabee, 0.9) were found in the AIGT+Soytide diet sample groups. Overall the distances within the various diet samples varied (**Figure 3**).

**Figure 3.**
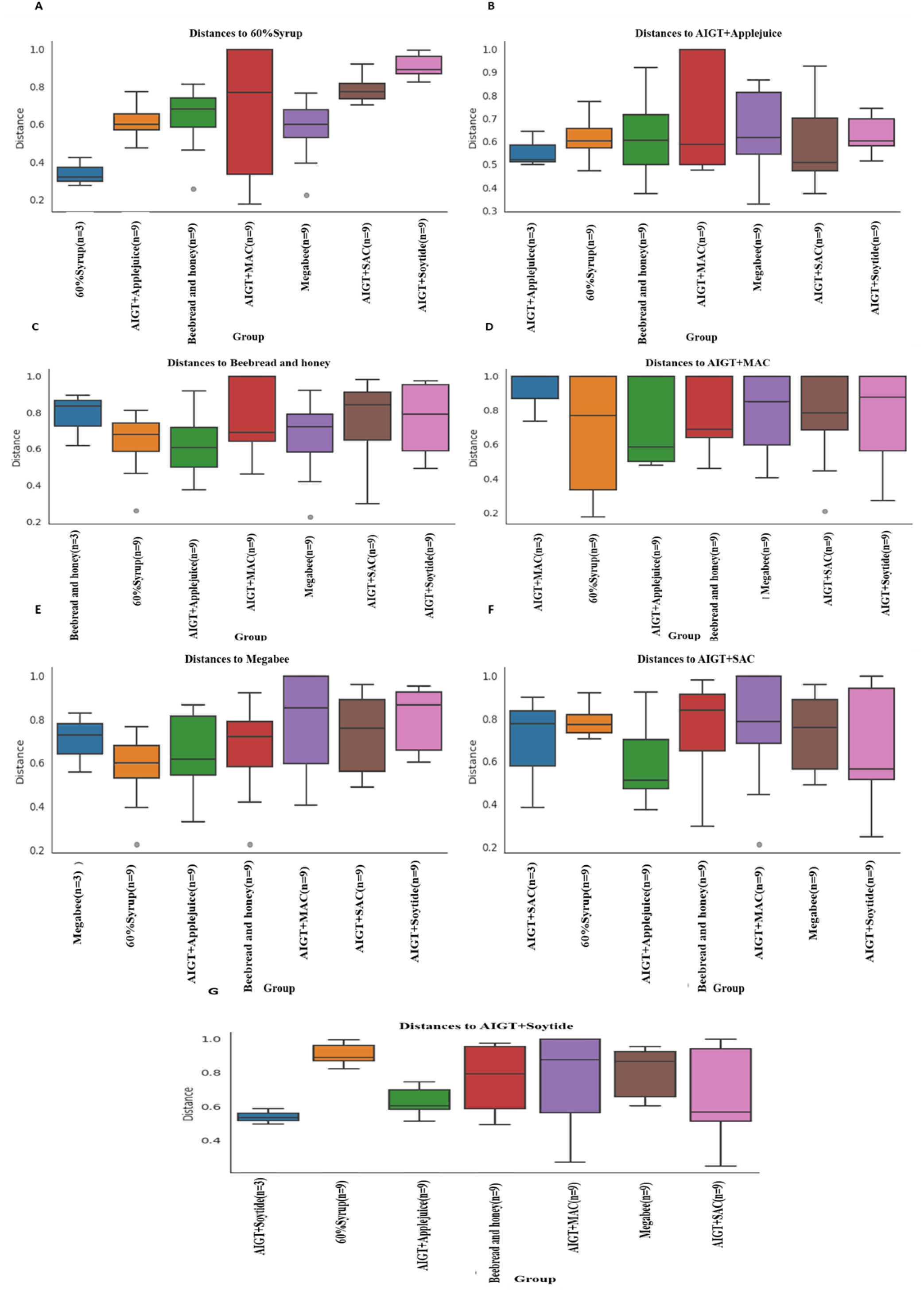
**Beta** Diversity group significance of samples. Beta diversity of the microbiome in seven different diet groups including AIGT+Apple juice, Beebread and honey, AIGT+MAC, Megabee, AIGT+SAC, AIGT+Soytide, and 60% Syrup was analyzed. The beta diversity; unweighted UniFrac distance to 60% Syrup **(A)**, AIGT+Apple juice **(B)**, Beebread and honey **(C)**, AIGT+MAC **(D)**, Megabee **(E)**, AIGT+SAC **(F)**, and AIGT+Soytide **(G)** of the microbiome in the gut of *Apis mellifera* exposed to these diets. N matches sample for each diet group; the 60% Syrup, AIGT+Apple juice, Beebread and honey, AIGT+MAC, Megabee, AIGT+SAC, and AIGT+Soytide have 9 (=3×3) respectively.

### 3.3 Gut Microbiota Taxonomy Diversity Based on Different Diets in Apis mellifera

The microbiome profile was analyzed based on different diet groups, and at different taxonomic stages including class, family, and genus level respectively (**Figure 4A-C**). At the class level, the most common bacteria, and widespread among the different diet groups were *Bacilli* followed by *Alphaproteobacteria,* and *Gammaproteobacteria* (**Figure 4A**). AIGT+SAC and AIGT+Soytide shared more similar microbiome compared to other diets at the class level. Even though *Gammaproteobacteria* was abundant in all the diet groups, however, it is abundant was relatively higher in the AIGT+MAC diet group. Similarly, *Alphaproteobacteria* showed higher abundance in the 60% Syrup diet group and less abundance in the AIGT+SAC, and AIGT+Soytide diet groups respectively. Also, at the family level, *Lactobacillaceae* remains a widely spread bacteria among the different diet groups (**Figure 4B**).

**Figure 4.**
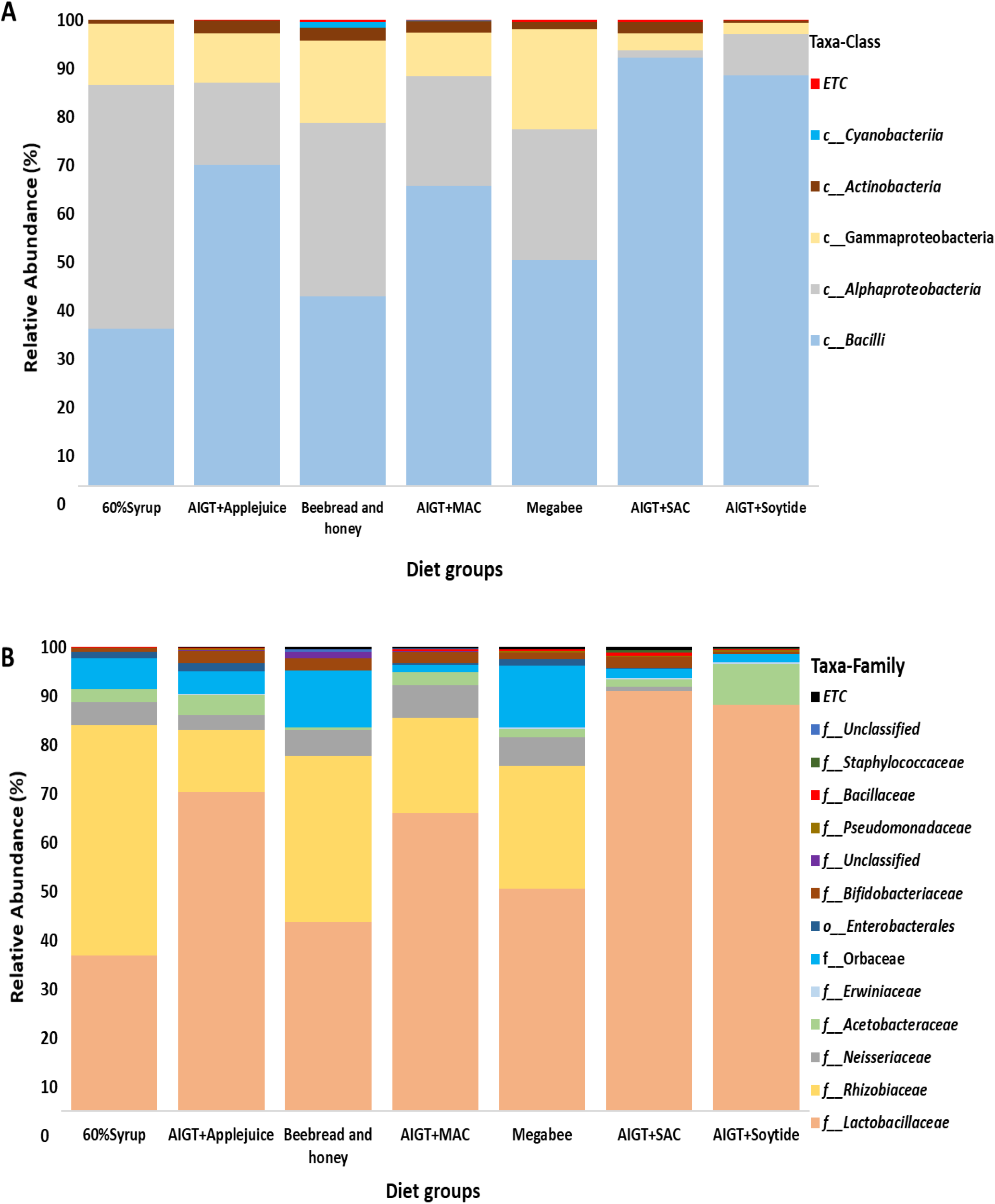
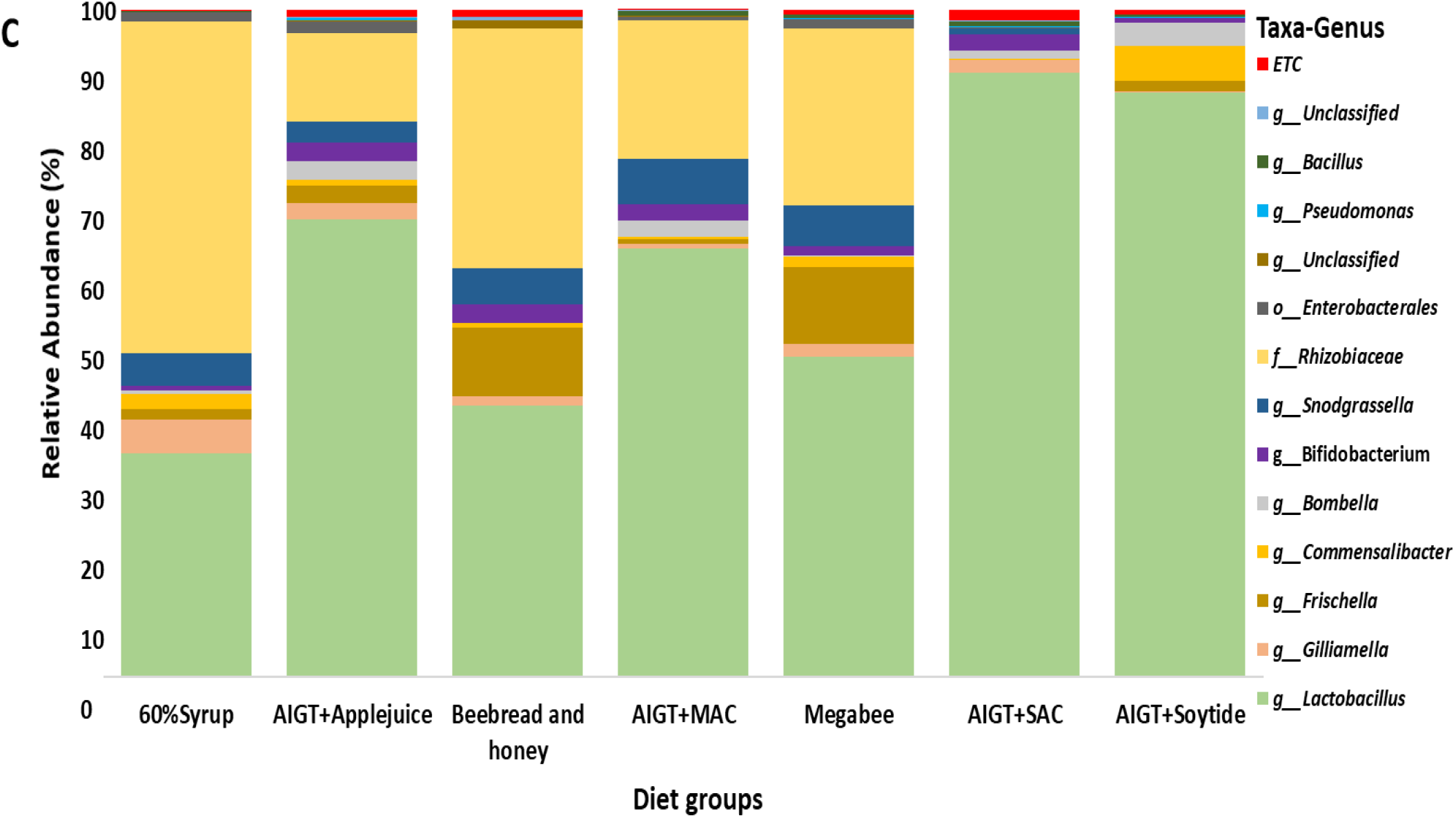
Microbiome profiles of *Apis mellifera* species using different diet compositions at Class **(A)**, Family **(B),** and Genus level **(C)** respectively. The bar height represents the average relative abundance of each taxa. The horizontal names represent the names of various diet groups. The microbiome composition was analyzed per each diet group including 60% Syrup, AIGT+Apple juice, Beebread and honey, AIGT+MAC, Megabee, AIGT+SAC, and AIGT+Soytide.

At the genus level also, data was analyzed to identify the relevant bacterial operational taxonomic units (OTUs) distributed for each diet group sample and among all the *Apis mellifera* samples. In total, 37 genera were found among all the diet groups. The most abundant genera/taxa include *Lactobacillus* (62%), followed by *Rhizobiaceae* (21%), *Snodgrassella* (4%), and *Erwiniaceae* (4%) among others. *Lactobacillus* not only occupies the highest abundance among the microbiome distribution, but also remains highly distributed among the diet groups in all the samples and equally dominated the microbiome abundance in AIGT+Soytide, AIGT+SAC, Megabee, AIGT+MAC, and AIGT+Apple juice diets groups, except in 60% Syrup, as well as Beebread and honey (**Figure 4C**). *Rhizobiaceae* dominated 60% Syrup, with significant abundance in Beebread and honey diet groups. It is important to highlight that, researchers do come across some unusual bacteria in the honey bee microbiome including the mentioned *Rhizobium,* which is well-known as nitrogen-fixing bacteria found in root nodules of legume plants [4]. These are likely *Gillamella* belonging to the order Rhizobium, yet are considered as part of the honey bee core microbiome [31].

Further, taking the diet groups together, *Lactobacillus* dominated the AIGT+Soytide, AIGT+SAC, AIGT+MAC, and AIGT+Apple juice. Also slightly dominated Megabee, and Beebread and honey, while *Rhizobiaceae* shows significant abundance at 60% Syrup (**Figure 4C**). *Frichella* is also widely distributed among all the diets, even though with less abundance in most of the diets except in the Beebread and honey, and Megabee diet groups. Similarly, *Bifidobacterium* distribution was observed in all the samples with almost similar abundance proportions. *Snodgrassella* was not found significantly distributed in the AIGT+Soytide diet group alone. *Bombella* abundance was also not significant in the Beebread and Honey diet group. *Commensalibacter* on the other hand was not also significantly found in the AIGT+SAC diet group but rather in all other groups (**Figure 4C**).

Further, at the genus level percentage of total reads per diet was analyzed and filtered out the microbiome distributions and proportions according to the diet groups (**Figure 5**). For the 60% Syrup diet, *Rhizobiaceae* was dominant with around (50%) followed by *Lactobacillus* with (34%), and *Gilliamella* with (5%). AIGT+Apple juice diet has *Lactobacillus* (69%) as the dominant genera followed by *Rhizobiaceae* (13%). The Beebread and honey diet shows the highest abundance rate of *Lactobacillus* (41%), followed by *Rhizobiaceae* (36%), and *Snodgrassella* (10%). AIGT+MAC diet group also shows *Lactobacillus* with the highest read (64%), followed by *Rhizobiaceae* (21%), then *Snodgrassella* (7%). Megabee diet consists of a large proportion of *Lactobacillus* (48%), followed by *Rhizobiaceae* (26%), and *Frischella* (11%). AIGT+SAC diet shows the highest percentage read of *Lactobacillus* (91%) among other diets, followed by *Bifidobacterium* (2%), and *Gilliamella* (2%). Lastly, the AIGT+Soytide diet group similar to other groups shows the highest abundance of *Lactobacillus* (88%), followed by *Commensalibacter* (5%), and *Bombella* (4%). Overall, among the 37 genera identified, only *Lactobacillus, Commensalibacter, Frischella, Bifidobacterium,* and *Gilliamella* taxa are present in all the seven different diet groups, and only *Lactobacillus* remain present in each sample among the total of 21 samples within the groups **(Table S4)**. Other bacteria such as *Prevotella* and *Clostridium* are found in only one sample each among 21 samples.

**Figure 5.**
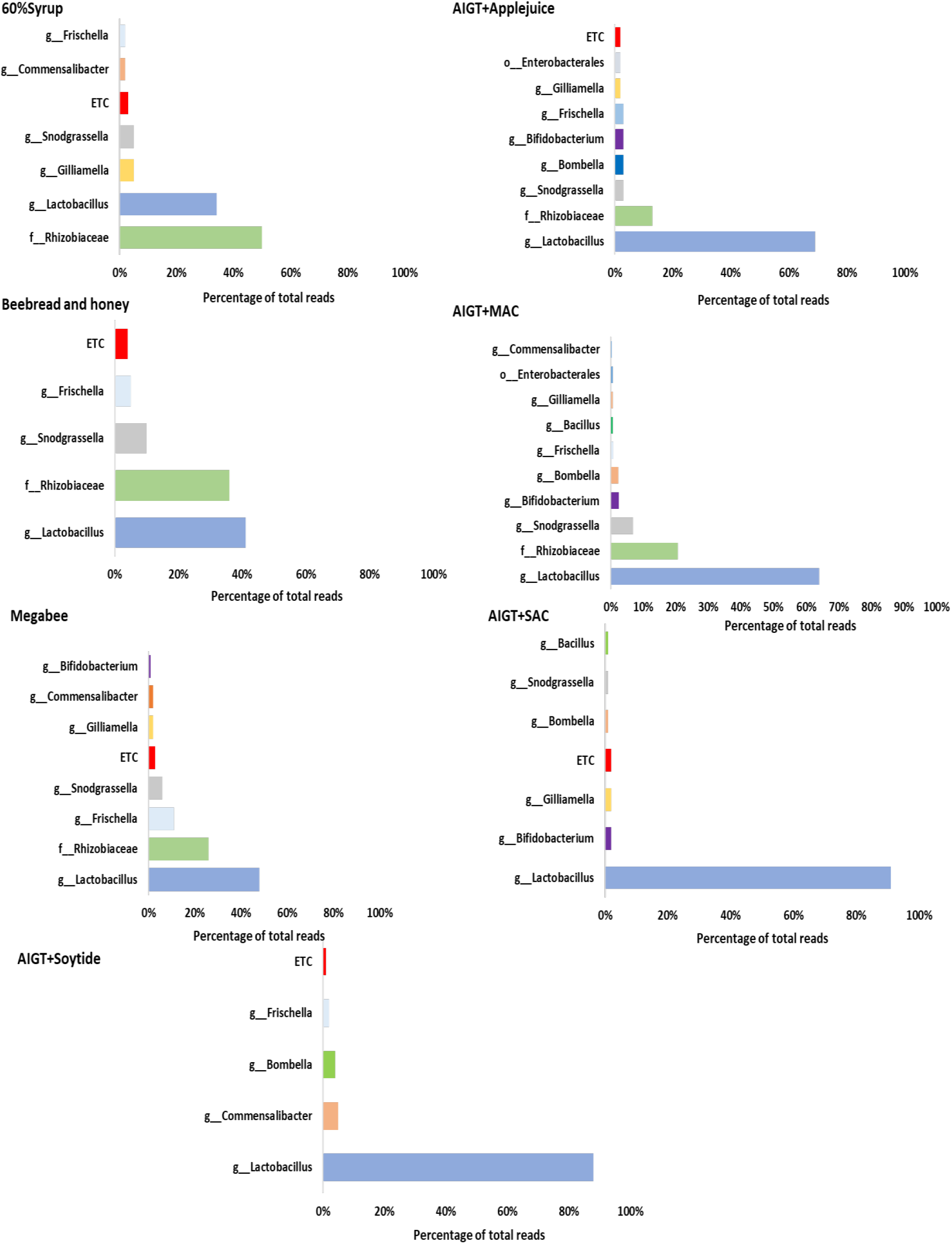
Microbiome profiles of *Apis mellifera* species using different diets at the genus level. The bar represents the percentage of total reads per diet group. The microbiome composition was analyzed per each diet group including 60% Syrup, AIGT+Apple juice, Beebread and honey, AIGT+MAC, Megabee, AIGT+SAC, and AIGT+Soytide. In the groups, *Lactobacillus* shows the highest reads among the diet groups, except in 60% Syrup which has *Rhizobiaceae* with the highest read.

### 3.4 Honey bee Health Condition Depends on Different Diets

To evaluate the health condition after feeding with various types of diets, we estimate the consumption, protein content in the head, and Vg expression level. During the five-day experiment, the consumption of diet by one honey bee was recorded daily, while taking into account the number of live honey bees each day. The cumulative consumption over the five days is presented in (**Figure 6A**). There are significant statistical differences between the diets (P < 0.05). AIGT+MAC (177.6 mg/bee) exhibited the largest consumption among the diets, followed by 60% Syrup (160.9 mg/bee), Beebread and honey (156.1 mg/bee), AIGT+Soytide (149.7 mg/bee), AIGT+Apple juice (146.0 mg/bee), Megabee (141.4 mg/bee), and AIGT+SAC (134.6 mg/bee), which showed the smallest consumption. In consumption, AIGT+MAC showed significantly higher than Megebee, Beebread and honey, and 60% Syrup. After 5 days of feeding the diets, protein content in the head was statistically significant between the diets (P < 0.05) in (**Figure 6B**). AIGT+SAC (418.4 µg/ml) showed the highest protein content in head among the diets, followed by AIGT+MAC (312.6 µg/ml), AIGT+Soytide (309.5 µg/ml), Megabee (290.2 µg/ml), AIGT+Apple juice (246.6 µg/ml), Beebread and honey (242.5 µg/ml), and 60% Syrup (223.8 µg/ml). The Vg expression level was statistically significant in diets (P < 0.05) in (**Figure 6C**). AIGT+Soytide (167.7) showed the highest level followed by AIGT+SAC (167.6), Beebread and honey (114.3), AIGT+Apple juice (70.2), Megabee (62.4), then AIGT+MAC (42.5). Among our developed diets, AIGT+Soytide and AIGT+SAC showed statistically higher Vg expression levels than Megabee. Additionally, AIGT+Soytide showed statistically higher Vg expression levels than Beebread and honey.

**Figure 6.**
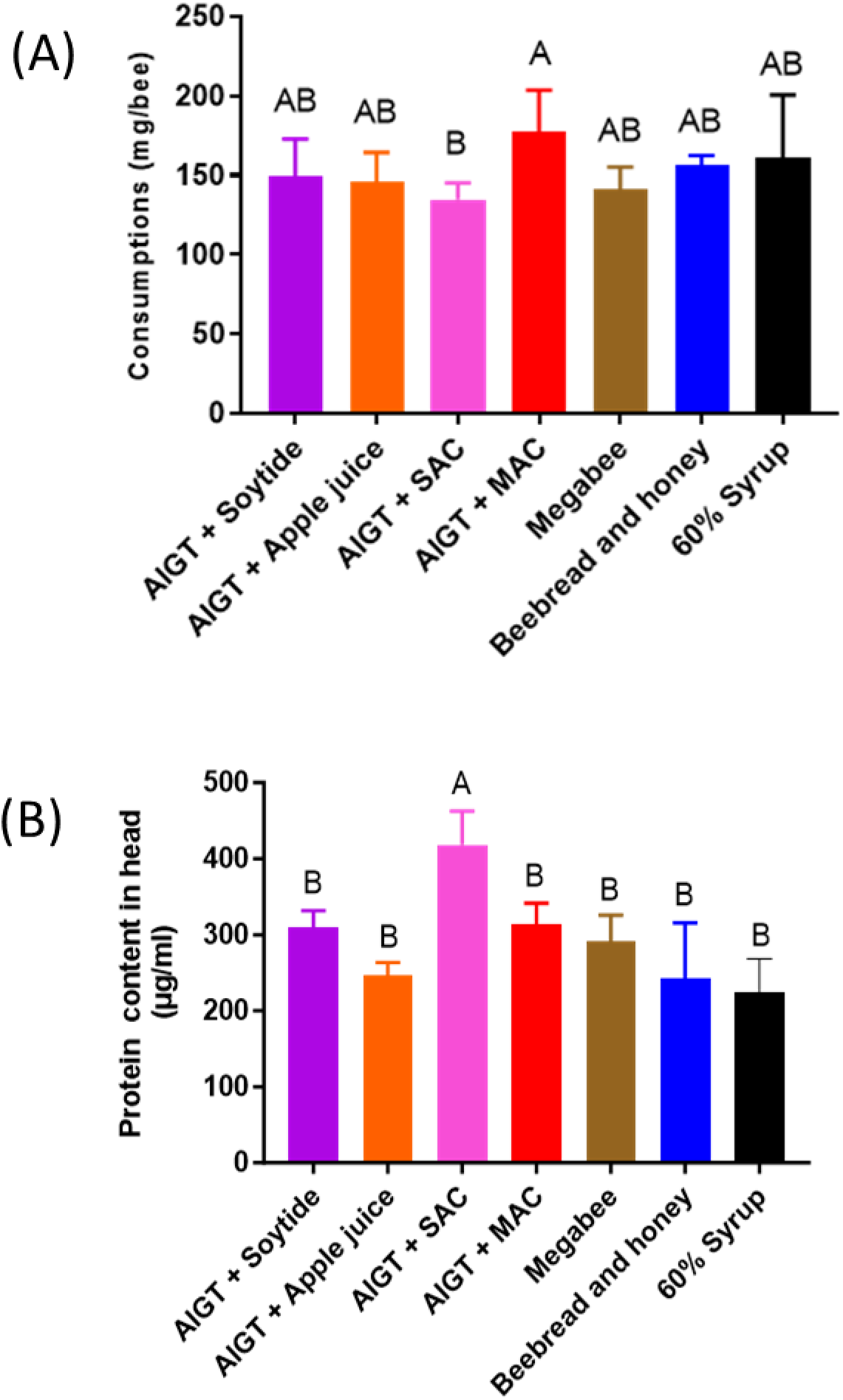
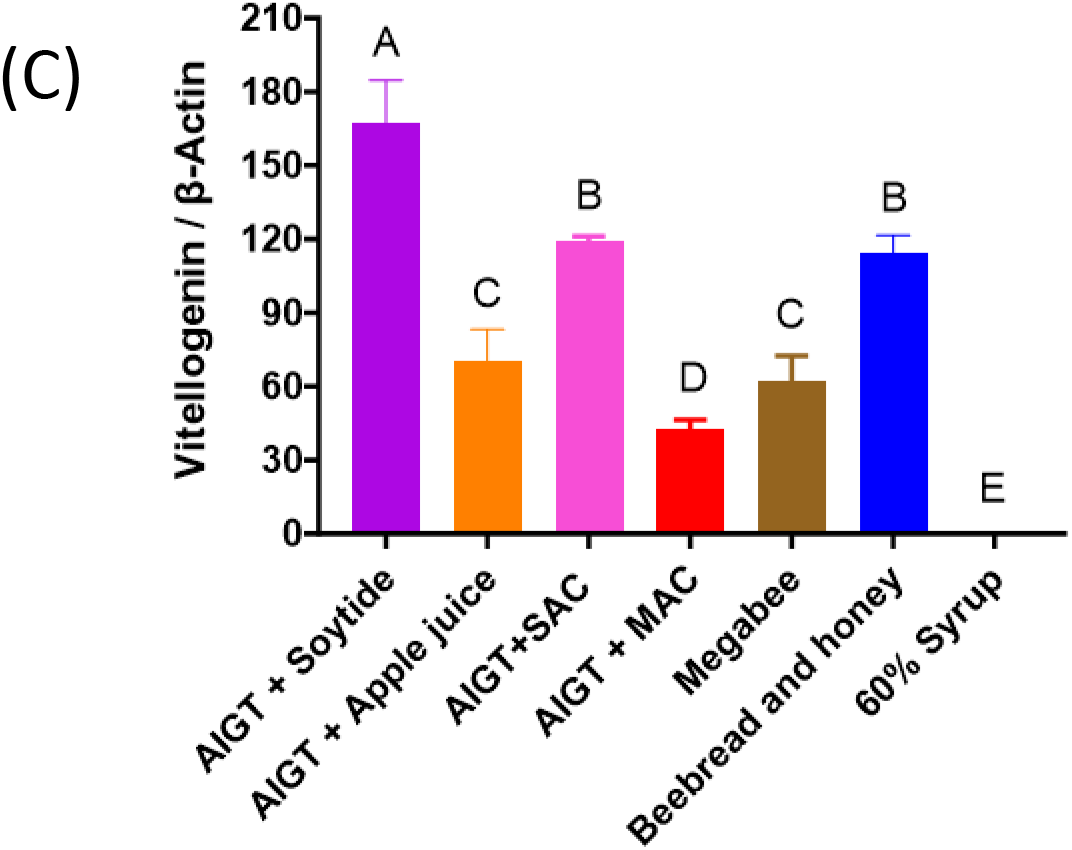
Bar plots showing the honey bee health effects based on the different diets from the cage experiment. (**A**) The diet consumption depends on the different diets. (**B**) Protein content in the head according to different diets. (**C**) The vitellogenin expression level of different kinds of diets. The means followed by different letters are significantly different according to Duncan’s multiple range comparisons (DMRTs) (*P* < 0.05).

## 4.0 Discussion

The importance of microbiomes is not limited to their interaction with their host but rather extended to their total compositions, and their metabolic and gene products related to the host environment and lifestyle including nutrition [5,32]. These together, influence the host’s biology, behavior, development, and immunity [33,34]. Communities of symbiotic microorganisms that colonize the gut play a significant role, especially against opportunistic microbes. However, diet diversity increases the number of symbionts which becomes a benefit for the host [35]. Considering this positive effect, we tested seven different diets and fed them to our model organism (*Apis mellifera*). This could help us to distinguish not only the best diet for nutritional quality but also the best diet that gives a healthy microbiome community against symbiotic microorganisms and or pathogens.

Diets with different compositions indirectly promote changes in the host-associated bacterial community [36]. Interestingly, from our findings, the alpha diversity measurements of the relative abundance of bacteria between the diet groups have shown that there are wide changes in microbiome compositions between the diet groups especially between Beebread and honey and Megabee as well as between AIGT+SAC and AIGT+Soytide. Other diet groups show a weak correlation between the microbiome of different diets. In the past, a relationship between the content of the insect’s diet and bacterial diversity linked with the host was observed in different insect species, such as *Plodia interpunctella; Blattella germanica, Drosophila* spp, and *Lymantria dispar* [37,38]. From our study using *Apis mellifera,* the bacterial communities associated with the different diet composition show less number of OTUs compared with OTU abundance reported in other insects [36]. This may be a result of close compositions among the different diets used except the 60% Syrup diet which has an entirely different composition with higher sugar content, and no protein. Our PCoA analysis is in line with that, considering the AIGT+Soytide microbiome from all three samples are closer to one another and distinct from other samples. In armyworm, diet source such as soybean was reported to be important in shaping the midgut bacterial communities [39], indicating the relevance of AIGT+Soytide as a good diet for the microbiome. Further, under natural conditions, increased diet diversity typically signifies a higher diversity of opportunistic microorganisms [40]. This is not the case in some studies, because the host received bacteria-free food [35]. However, similar to our finding, there is higher diversity among some of the diet groups after comparing the alpha and beta diversity.

The overall microbial relative abundance was analyzed and compared between the diet groups showing a wide spectrum of bacterial distributions among the groups. The dominant microbiota present in the guts of honey bees were Proteobacteria, Firmicutes (*Bacilli, Lactobacillus*), Actinobacteria (*Bifidobacterium*), and Bacteroides. These taxa are consistence with those found in a recent study [41,42]. The mentioned studies suggest that the dominant groups of bacteria in the honey bee guts may hardly changed by external factors such as temperature, seasons, toxins, and even nutrients such as sugar [43]. However, in our current findings, the relative abundance (RA) of the dominant microbiota changed with different diet types. For example, *Lactobacillus* is the most abundant genera in the AIGT+Soytide diet followed by *Commensalibacter* and *Bombella.* On the other hand, *Rhizobiaceae* is the most abundant genera in the 60% Syrup diet group followed by *Lactobacillus* and *Gilliamella.* This may be related to the diet compositions which can influence the microbiome abundance. For example, market high-fructose syrup inevitably contained proteins and polysaccharides that were difficult to digest resulting in more feces in the hindgut of honey bees [44]. This provides changes in the microbiome in favor of the honey bee intestinal self-regulation system with varied dietary components and adapts to environmental changes.

At the genus level, not only in the AIGT+SAC diet group, *Lactobacillus* (lactic acid-producing bacteria) family members show higher abundance in most of the other diet groups. Lactic acid bacteria (LAB) are found as a typical inhabitant in the insect gut such as cockroaches [45], termites [46], and honey bees [47]. The biological role of LAB especially concerning insect physiology is not fully elucidated. In the honey bee, LAB plays a protective role against pathogens [48], and support respiratory requirement in cockroaches [45]. Therefore, a higher abundance of the LAB in different diet groups can be considered as a marker for diet quality. From our findings, at the genus level, *Lactobacillus* bacteria dominate the AIGT+Soytide, and AIGT+SAC diet groups with RA>80% respectively. While 60% Syrup, and Beebread and honey diet groups show a poor abundance of *Lactobacillus* with RA <50% as compared to other diet groups. The identification of *Lactobacillus* in the microbiota control by these diets represents an interesting discovery because of its important role in insect biology [49]. Recently, *Bifidobacterium* is also considered as a LAB [49]. Both LAB and acetic acid bacteria (AAB) are common in the honey bee gut [50,51]. The AAB have some potentially beneficial effects on the host [51]. While the LAB increase honey bee immunity and protects the host from bacteria, yeast, and pathogens [52]. Host susceptibility to disease increases when the abundance of LAB and AAB is low [53]. Added to that, beekeepers use lactic, and acetic acids to protect the honey bees against pathogens [54], indicating the vital role of these bacteria against the honey bee pathogens. In our current analysis, AIGT+Soytide, and AIGT+SAC diets increased the relative abundance of Firmicutes (*Lactobacillus*), and Actinobacteria (*Bifidobacterium*) in the gut, most of which are beneficial bacteria compared with other diets especially sugary diet (60% Syrup). On the other hand, a decrease in the RA of Alphaproteobacteria, and Gammaproteobacteria which contain numerous medically important bacteria including the pathogenes was observed from the same diet groups (AIGT+SAC and AIGT+Soytide diets). The genera *Commensalibacter* and *Bombella* are also AAB belonging to the *Acetobacteriaceae* family and have the ability to colonize honey bee-associated environments such as honeycombs and beebread as well as honey bee gut [10,55]. *Commensalibacter* bacteria have been found in and isolated from the intestines of various insects that feed on high carbohydrate diets including the honey bees (*Apis mellifera*, *Apis florea,* and *Apis dorsata*) [56,57]. Several reports suggest that *Commensalibacter* are associated with the health of their respective insect hosts. For example, *C. intestini* was reported to be involved in modulating Drosophila immunity [51], highlighting it is importance in insect health quality. From our analysis, the highest abundance of the *Acetobacteriaceae* family (*Commensalibacter,* and *Bombella*) was observed in the AIGT+Soytide diet group. From cage experiment results, the Vg expression level which is considered as a honey bee health biomarker [58], showed the highest expression in the AIGT+Soytide group among the other diet groups. Vg affects worker sucrose response, foraging initiation, foraging choice, caste differentiation process, protection from oxidative and immune attacks, and longevity [59–61]. Therefore, these results indicate the significance of this diet in honey bee health. To further fish out the best diet for honey bee health, we identified that *Rhizobiaceae* RA is significant in most of the diet groups except in AIGT+SAC, and AIGT+Soytide. Interestingly, high levels of *Rhizobiaceae* have previously been associated with honey bees fed only with sugar syrup [62]. Not surprisingly, from our findings, *Rhizobiaceae* RA was highest (50%) in 60% Syrup diets that are highly rich in sugar components as compared to all other diets including the control (Megabee). *Rhizobiaceae* abundance was influenced by tau-fluvalinate exposure [63], highlighting its response not only to sugar but also to xenobiotic stress.

Considering the influence of different diet groups on honey bee microbiota, the analysis of individual honey bee health status also aligns with the microbiome findings. This is observed by exploring the diet-microbiome relationship for honey bee health through the expression level of Vg. From the present study findings, the AIGT+Soytide group shows the highest expression level of Vg, followed by AIGT+SAC, and the Beebread and honey diet groups. This indicates the significance of these diets in honey bee health. Additionally, these groups are characterized by a high abundance of *Lactobacillus*, *Bifidobacteria, Snodgrasella,* and *Frischella* which have been identified as markers of honey bee health [64]. This indicates the significance of these diets in promoting honey bee health. Conversely, 60% Syrup shows the lowest Vg expression level. In microbiome results, *Rhizobiaceae,* a marker for poor honey bee health [65], has the highest RA in the 60% Syrup group, indicating that this diet may not contribute to the overall health of honey bees. Moreover, the honey bee group fed with AIGT+SAC shows the highest protein content in the head. The Protein content in the head and the development of the hypopharyngeal gland are linked to the secretion of royal jelly, which serves as nourishment for both larvae and queens [26]. AIGT+SAC group exhibits statistically significantly higher Vg expression level than Megabee. This suggests that AIGT+SAC could contribute to the promotion of individual honey bee health and potentially impact the development of the honey bee colony. The excessive dominance of *Lactobacillus* (LAB) and the significant abundance of *Bifidobacteria* (LAB) alongside other AAB might have influenced the absorption of protein content in the *Apis mellifera* head through the gut-brain-axis, thereby being associated with honey bee health. It is noteworthy that honey bee health correlates with consumed nutrition [66]. Further, AIGT+SAC and AIGT+Soytide display high-efficiency rates in terms of individual honey bee health. Overall, our definitive findings indicate that different diets have varying effects on honey bee microbiome health status. Therefore, this suggests a link connecting diet, honey bee health, and microbiome.

## 5.0 Conclusions

The present study demonstrated the impact of different diets on the microbiome community of *Apis mellifera* and its consequential effect on honey bee health. Comparative genomic analysis revealed that the AIGT+SAC and AIGT+Soytide diet groups exhibit a higher abundance of LAB and AAB, including *Lactobacillus, Bifidobacterium, Commensalibacter,* and *Bombella.* These microbial indicators are associated with improved honey bee health and enhanced immunity when present in the appropriate proportions. The ability of LAB to increase honey bee immunity and protect the host from pathogens provides valuable insights for identifying high-quality diets. The study establishes a correlation between diet quality, individual honey bee performance as indicated by the expression level of Vg, and the protein content in the head. Consequently, we propose that diet compositions such as AIGT+SAC and AIGT+Soytide are likely to offer better nutritional quality, shaping the microbiome community into a healthier state. Furthermore, it is recommended that the relationship between diet-microbiota and honey bee competency be further explored through colony-level experiments rather than cage experiments. This is crucial as microbiota abundance can be influenced by environmental factors.

## Supplementary materials

The following supporting information can be downloaded at: www.mdpi.com/xxx/s1, **Table S1**: Primer information for DWV, Vg, and ß-actin; **Table S2**: Primer for amplifying V3-V4 region; **Table S3**: The table indicates the summary of the Next-generation sequencing (NGS) analysis; **Table S4**: Showing the commonly distributed taxa among the diet groups.

## Author contributions

H.W.K., H.J.K designed research; H.J.K, A.Y.M., and J.H.L., performed research; A.Y.M., J.H.L, and H.J.K, analyzed data; A.Y.M., Writing-original draft of the paper. H.J.K. and F.O. performed sampling. H.W.K. A.Y.M., H.J.K, J.H.L., and M.B. Writing-review and editing. All authors reviewed the manuscript. The author(s) read and approved the final manuscript.

## Acknowledgments

This work was carried out with the support of Cooperative Research Program for Agriculture Science & Technology Development (Project No. PJ015755022023) and the Priority Research Centers Program through the National Research Foundation of Korea (NRF) funded by the Ministry of Education (2020R1A6A1A03041954); and Research Assistance Program (2021) in the Incheon National University.

